# Losartan promotes cell survival following SARS-CoV-2 infection *in vitro*

**DOI:** 10.1101/2020.12.27.424507

**Authors:** Reza Nejat, Ahmad Shahir Sadr, Brendan T. Freitas, Jackelyn Crabtree, Scott D. Pegan, Ralph A. Tripp, David J. Najafi

## Abstract

**Introduction:** Coronavirus disease 2019 (COVID-19) can be associated with mortality and high morbidity worldwide. There is an extensive effort to control infection and disease caused by SARS-CoV-2. This study addressed the hypothesis that angiotensin II type I receptor blocker, Losartan, may restrict pathogenesis caused by SARS-CoV-2 by decreasing viral-induced cytopathological changes by blocking angiotensin II type 1 receptor (AT1R), thus reducing the affinity of the virus for ACE2, and inhibiting papain-like protease of the virus.

**Method:** Losartan inhibitory effect on deubiquitination and deISGylation properties of papain-like protease was investigated using a fluorescence method and gel shift analysis determining its inhibitory effects.

The inhibitory effect of Losartan on SARS-CoV-2 cell replication was investigated both when losartan was added to the cell culture 1 hour before (pre-infection group) and 1 hour after (post-infection group) SARS-CoV-2 infection of Vero E6 cells.

**Results:** Losartan treatment of Vero E6 cells prior to and after SARS-CoV-2 infection reduced SARS-CoV-2 replication by 80% and 70% respectively. Losartan was not a strong deubiquitinase and deISGylase inhibitor of PLpro.

**Conclusion:** Losartan added pre- and post-infection to the Vero E6 cell culture significantly prevents cell destruction and replication by SARS-CoV2. Losartan has low side-effects, is readily available, and can be produced at high levels globally, all features of a promising drug in treatment of COVID-19 if validated by clinical trials.

## Introduction

The COVID-19 pandemic having with high morbidity and substantial mortality rates has created a global health and economic crisis since December of 2019. It is caused by severe acute respiratory syndrome coronavirus 2 (SARS-CoV-2). SARS-CoV-2 is an enveloped non-segmented single-stranded positive sense RNA virus belonging to the beta-genus of the Orthocoronavirinae subfamily in the Coronaviridae family which includes SARS-CoV and Middle East Respiratory Syndrome CoV (MERS-CoV) [1, 2]. Since its emergence in Wuhan, China, SARS-CoV-2 has infected over 76 million and caused 1.7 million deaths globally according to the World Health Organization (WHO) status report on 23 of December 2020.

As a multi-organ disease, COVID-19 involves lungs, cardiovascular system, gastrointestinal tract and the central nervous system. This virus at the genomic level has 79% homology with SARS-CoV [1–4]. There is currently no effective antiviral drug and currently developed vaccines are not expected to be administered worldwide by the end of 2021. Discovery of a drug that is globally available with antiviral properties and minimum side-effects would significantly impact the management of the pandemic.

SARS-CoV-2, like SARS-CoV, uses the ACE2 receptor, a member of renin-angiotensin system, to enter host cells [5]. Binding of the virus spike (S protein) protein, through its receptor binding domain (RBD) in S1 component to ACE2 mediates the cell entry [6]. Inhibition of this process halts infection and disease pathogenesis of COVID-19.

Entry of the virus into the host cell initiates its replirrecation. Translation of two open reading frames, (ORF1a/ORF1b) produces two replicase polyproteins (pp1a/pp1ab) [7, 8]. These polyproteins each contain non-structural proteins (nsps) involved in viral replication. Cleavage of these replicases releases 16 nsps through an autoproteolyzing process. Sharing the first 11 nsps, pp1a and pp1ab release nsp1-11 and nsp1-16, respectively [9, 10]. Two cysteine proteases, papain-like protease (PLpro) and 3Chemotrypsin-like proteinase (3CLpro or Mpro), contribute to this proteolysis. PLpro is located in the sequence of nsp3 which separates nsp1/nsp2, nsp2/nsp3 and nsp3/nsp4 while Mpro, embedded in nsp5, separates the nsp4-11 of pp1a and nsp4-16 of pp1ab [10, 11]. Mpro has been shown to reduce induction of type I interferon [12]. In addition, PLpro of SARS-CoV-2, with 83% homology with its counterpart in SARS-CoV, has deubiquinating and deISGylating potential with differing substrates. PLpro of SARS-CoV and SARS-CoV-2 show predominance in cleaving ubiquitin-like interferon-stimulated gene 15 (ISG15) proteins over poly-ubiquitin chains. However, SARS-CoV cleaves ubiquitin more readily than SARS-CoV-2 [13, 14]. These properties modify and suppress the repertoire of host cell proteins and mediators such as IFN-β, ISG15 protein, IRF3, TLR3, TLR4, TLR7 and STING-mediated pathway engaged in innate immunity [14–21]. It is implied that both PLpro and Mpro influence the immunogenic molecular pathways which allows SARS-CoV-2 to evade recognition by innate immunity system which prolongs its incubation period in host cells.

PLpro, of SARS-CoV and SARS-CoV-2, cleaves nsp1, nsp2 and nsp3 at the **L**X**GG** sequence of their C-terminals. The **L**X**GG** is present in the C-terminus of ubiquitin and ISG15protein [22–24]. Therefore, iso- and endo-peptidase action of PLpro on suppression of SARS-Cov-2 is a desirable target to suppress its pathogenesis [22]. Inhibition of PLpro deubiquitinating and deISGylating action may prompt the innate immunity against the virus.

In the early months of 2020, angiotensin receptor blockers (ARBs) were noted to possibly provide protection against COVID-19 [25–27]. Stopping the use of ARBs was recommended in the early phase of the pandemic due to the concern over possible increase in viral load [28, 29]. Recent studies have revealed safety of ARBs in patients with COVID-19 [30, 31]. Importantly, a recent study proposed that Losartan, an ARB, could ameliorate complications, especially of acute respiratory distress syndrome (ARDS) associated with COVID-19 [27]. In a previous study, molecular dynamic (MD) analysis showed that Losartan could change the three-dimensional structure of ACE2, preventing establishment of strong bonds between RBD residues of the virus and the residues in ACE2 [27], and subsequently decrease the affinity of the virus to its receptor. Furthermore, the MD analysis showed that Losartan could change the conformational shape of PLpro by occupying the place for inhibitors of PLpro with high docking energy (Fig.6) [27]. This implies that Losartan may act as a PLpro inhibitor.

PLpro and PLP2 of other Coronaviruses have been shown to have a substantial impact on the pathogenesis of CoVs [32–35]. PLpro inhibitors in SARS-CoV-2, have shown to inhibit its replication through suppressing cleavage of nsps [36, 37]. Moreover, these inhibitors prevent PLpro from acting as a deubiquitinase (DUB) or deISGylase, the two properties that enable the virus to evade innate immunity [13, 14]. In addition, as ubiquitins are involved in normal regulatory cellular pathways, deubiquitinating activity of PLpro modifies the host cell environment promoting viral replication [38]. Further work on discoveries of noncovalent inhibitors of PLpro in 2008, which showed them to inhibit viral replication, was aborted soon [14, 39]. Here, we report the very weak inhibitory effect of Losartan against deubiquitinase and deISGylase properties of SARS-CoV-2 PLpro. In this *in vitro* study of Vero E6 cells, we show that adding Losartan pre- and post-infection with SARS-CoV2 considerably prevents viral replication.

## Results

### Deubiquitinase and deISGylase inhibitory activity of Losartan on SARS-CoV-2 PLpro

To characterize its inhibitory effects on SARS-CoV-2 PLpro, Losartan was incubated at various concentrations with SARS-CoV-2 PLpro and a peptide substrate containing the last five consensus amino acids of ubiquitin (Ub) and interferon stimulated gene product 15 (ISG15) in conjugation with a C-terminal 7-amido-4-methylcoumarin (AMC) fluorogenic reporter group (peptide-AMC). These assays revealed IC_50_ value of 1200±61μM for Losartan against peptide-AMC (Table 1).

**Table 1.** Percentile of inhibition and IC_50_ of cleavage of peptide-AMC, percentile of inhibition of ISG15-AMC and Ub-AMC in the presence of 200μM Losartan. IC_50_ of Losartan for cleavage of peptide-AMC is 1200±61 μM.

| | Peptide<br>(200$\mu M$) | Peptide $IC_{50}$<br>($\mu M$) | ISG15<br>(200$\mu M$) | Ub (200$\mu M$) |
| --- | --- | --- | --- | --- |
| Losartan | 35.1% ± 2.7 | 1200 ± 61 | 6.9% ± 3.4 | 2.3% ± 1.4 |

ISG15 is the preferred substrate of SARS-CoV-2 PLpro [14]. To test the inhibitory effect, Losartan was tested against ISG15-AMC and peptide-AMC. At 200μM, Losartan is less effective at inhibiting cleavage of ISG15-AMC than against peptide-AMC. SARS-CoV-2 PLpro has a strong preference for ISG15 over peptide as a substrate displacing competitive inhibitors with lower affinity for ISG15. These data suggest that Losartan, along with its structure, may be interacting with the P3/P4 pocket of the PLpro [40]. This pocket is adjacent to the active site and facilitates cleavage of Ubl substrates by binding the C-terminal leucine and arginine of the RLRGG motif [41]. Other small molecule PLpro inhibitors of similar size have been shown to bind in this pocket in both SARS-CoV and SARS-CoV-2 [39].

### Efficacy of Losartan in Inhibiting SARS-CoV-2 PLpro

To further explore the relationship between substrate affinity and inhibitor efficacy, Losartan was tested against Ub-AMC at 200μM. SARS-CoV-2 PLpro has a strong preference for ISG15 over Ub as a substrate [14], where the catalytic efficiency of the PLpro for Ub is approximately one-tenth that of ISG15 but more than 250 times that of the peptide-AMC. Therefore, competitive non-covalent inhibitors limit Ub-AMC cleavage at a rate somewhere between those of ISG15-AMC and peptide-AMC. Losartan had an inhibition rate of 2.3% when tested against Ub-AMC (table 1). Inhibition rate of Losartan was 6.9% against ISG15 cleavage (table 1). These differences between inhibitory rates of Losartan against ISG15-AMC and Ub-AMC are unlikely to have a qualitative difference in a biological system. However, to confirm this, molecular weight shift assays were performed with more biologically accurate substrates (Fig. 1).

**Fig. 1.**
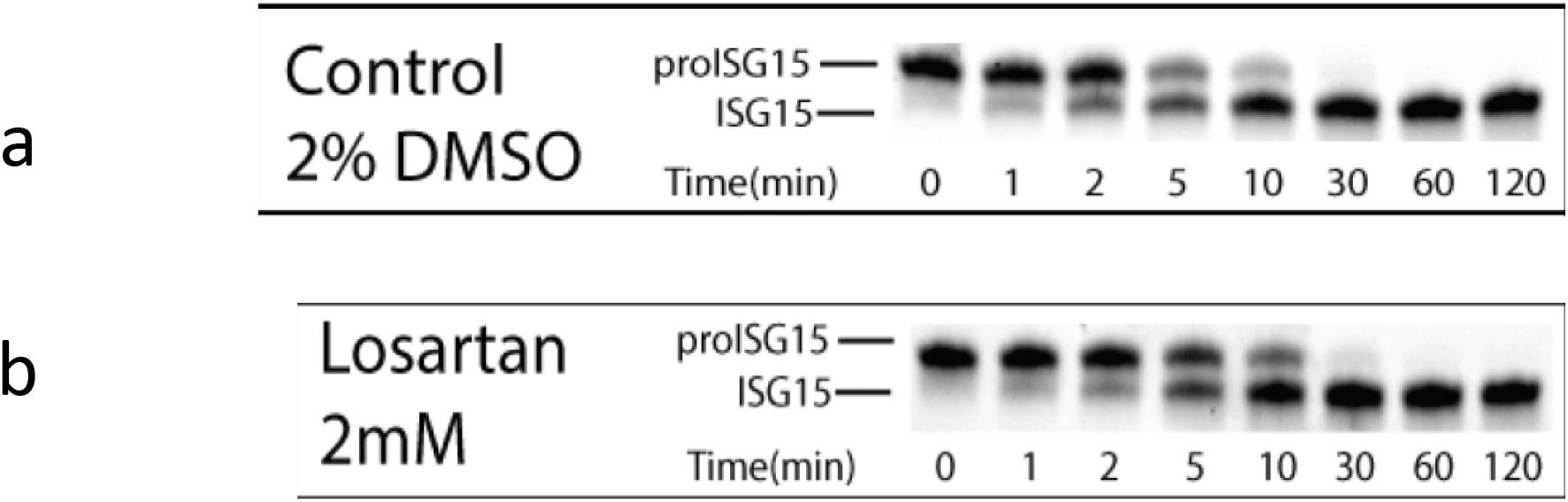
Gel shift analysis. Inhibitory activity of Losartan on deISGylation of PLpro. (a) deISGylation of PLpro on proISG15 in 10 μM of proISG run against 20 nM of SARS-CoV-2 PLpro, at 37°C, over 2 hours with samples taken at the indicated intervals with Gel cleavage assay visualized by Commassie Blue Staining. Strong PLpro deISGylation is noted. (b) Losartan, at 2mM concentration added to proISG15 and PLpro solution, demonstrates insignificant inhibitory effect on PLpro deISGylation between 5 and 30 minutes.

### Effect of Losartan on Tetra-Ub Deubiqutination

Ubl substrates with AMC tags at their C-terminal glycine are not entirely reflective of natural substrates. PLpros have been observed to prefer cleaving poly-Ub chains over mono-Ub substrates [42, 43]. To determine if the inhibition values derived from AMC cleavage assays are reflective of a qualitative change in DUB and deISGylase activity, Losartan was tested against human proISG15 and K48 linked tetra-Ub. As expected, Losartan at 2 mM shows a small deISGylase activity relative to the control (Fig. 1). However, Losartan at 2.25 mM almost completely eliminated DUB activity (Fig.2).

**Fig. 2.**
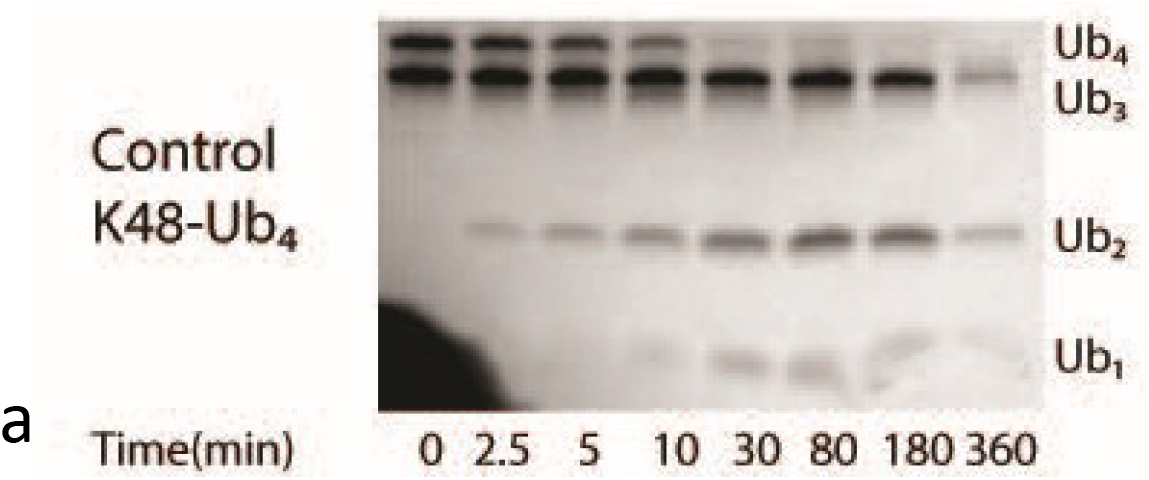
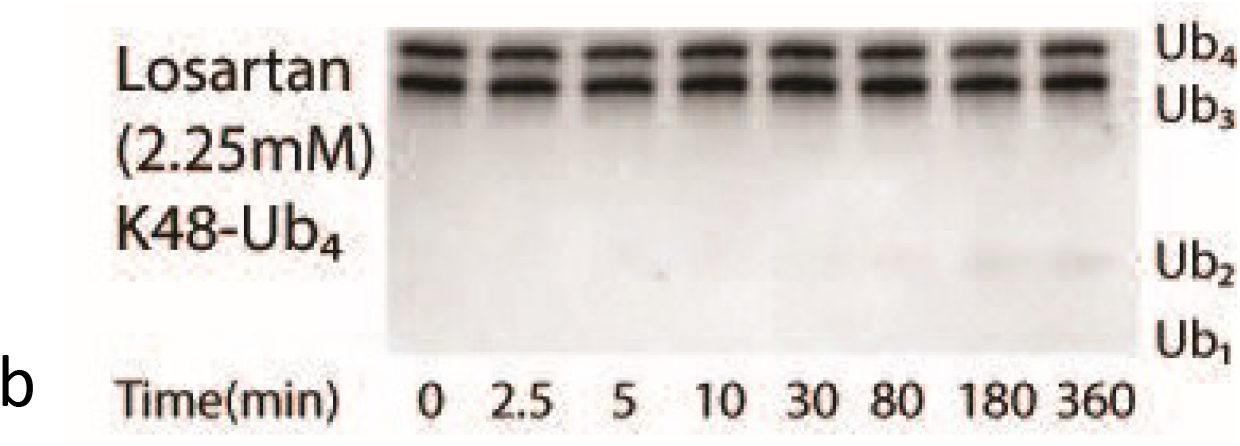
Inhibitory effect of Losartan on deubiquitination activity of PLpro on K48-Ub_4_ (a) At the mentioned intervals over 6 hours, multiple ten μL samples of Lys48 linked tetra Ub at 13.65 μM with 23 nM SARS-CoV-2 PLpro in AMC buffer at 37 °C were taken and heat-shocked at 98 °C for 5 minutes. Gel cleavage assay visualized by Commassie Blue Staining shows PLpro’s cleavage of Ub_4_ to mostly DUB and some monoUb. (b) Addition of Losartan, at 2.25 mM concentration, demonstrates a significant inhibitory effect on PLpro deUbiquitination of Ub_4_.

### Antiviral Activity of Losartan in Cell Culture

Adding Losartan to Vero E6 cell cultures showed that dose-dependently (0-100uM concentrations), it could significantly (p<0.001) reduce cell destruction and cell replication as noted in 6-well and 96-well assays, respectively (Figures 3 and 4). This result was achieved in both pre- and post-infection groups in both of 6-well and 96-well setting. In 6-well assay, Losartan increasing concentrations reduced the number of plaques up to 70% and in 96-well assay the number of infected cells by up to 80% (p<0.001). The half maximal effective concentration (EC_50_) of Losartan in 96-well experiment significantly (p<0.001) decreased from 40.81 μM in post-infection group to 13.65 μM in the pre-infection group (Fig. 5).

**Fig. 3.**
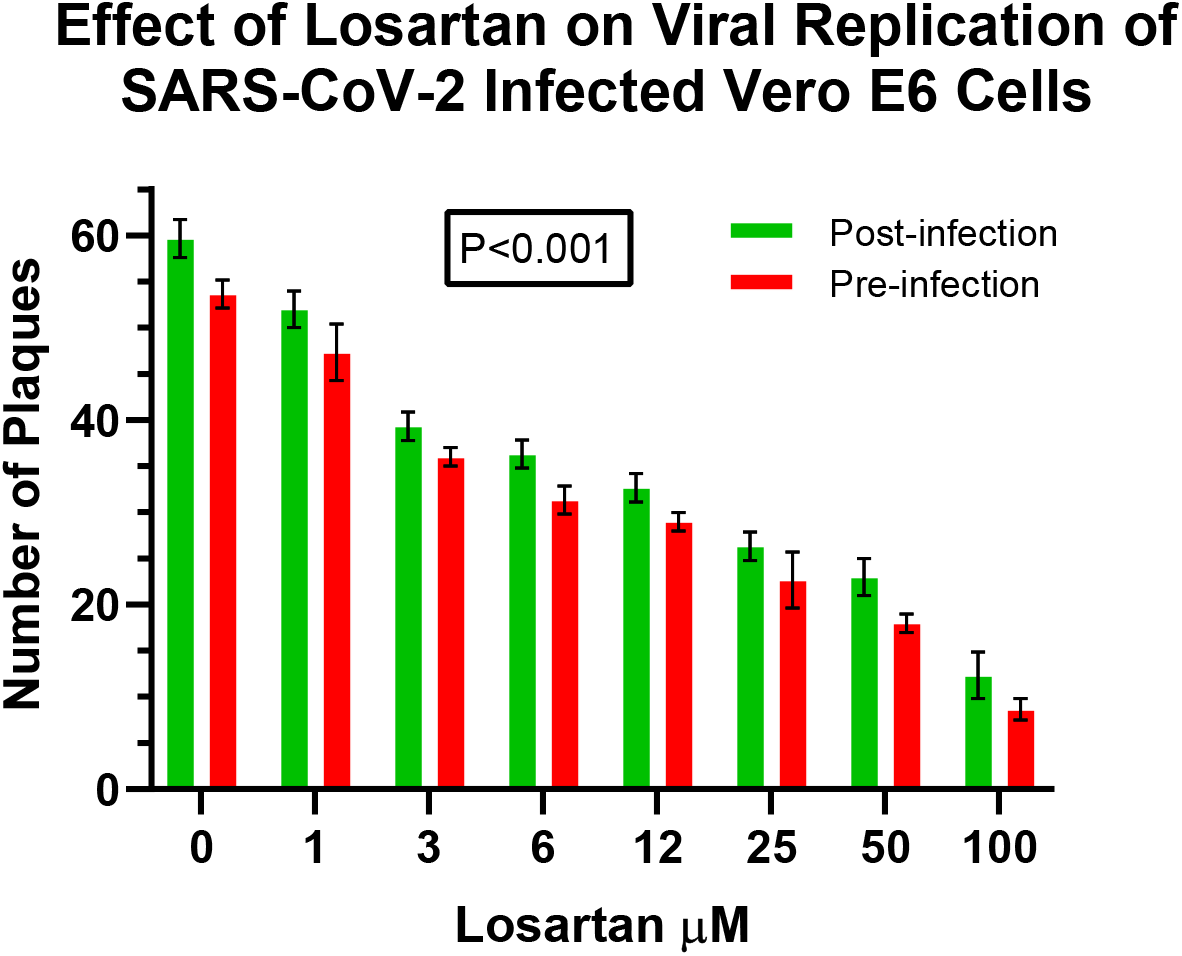
Losartan, in a dose dependent manner, decreases virus induced cell destruction significantly (p<0.001with Wilcoxon test), i.e. the number of plaques, by better than 80%. Pre-infection and post-infection groups indicate that Losartan was added one hour before (pre-infection) and after the cells were infected (post-infection) with SARS-CoV-2, respectively.

**Fig. 4.**
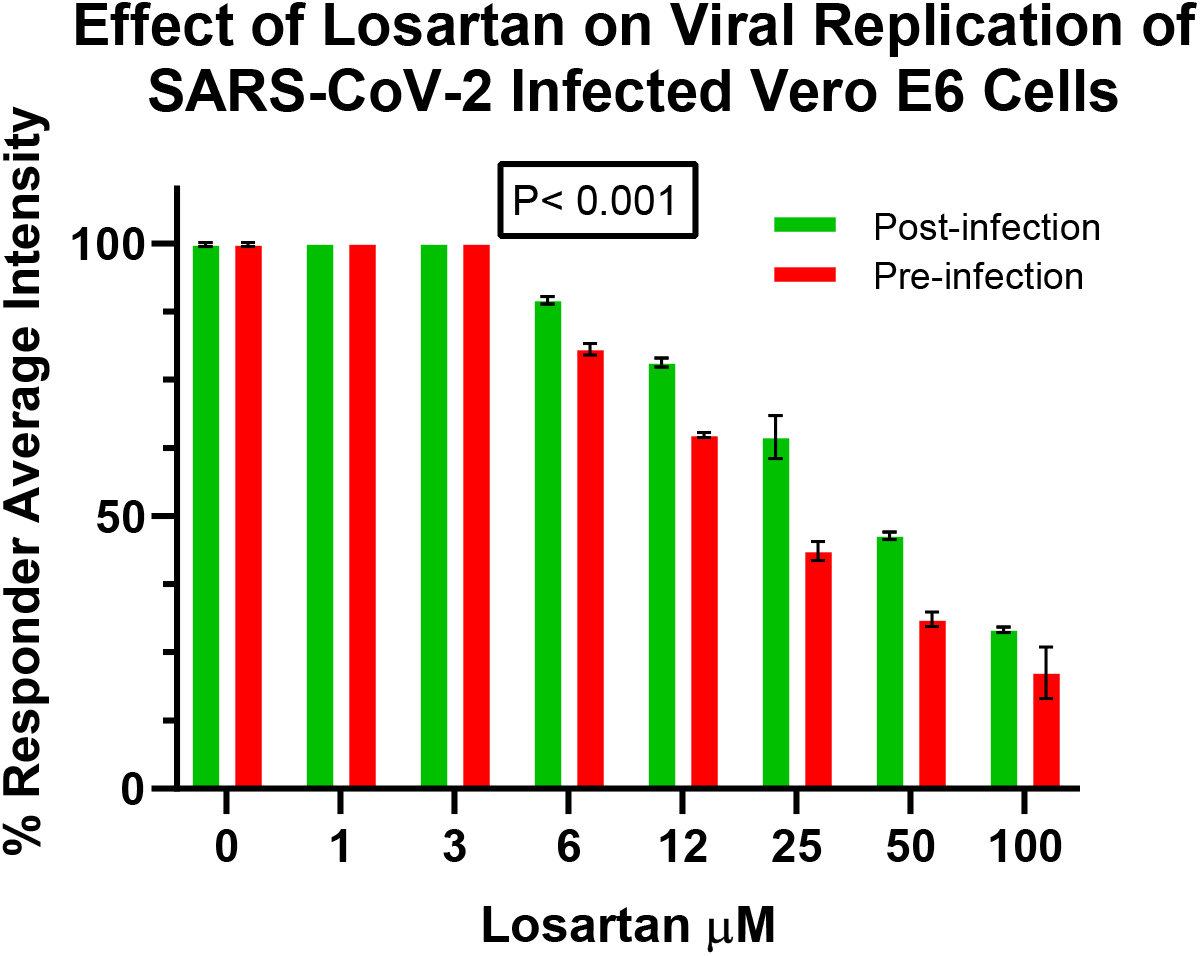
Losartan, in a dose dependent manner, decreases the percentage of infected cells in pre-infection group more significantly than post-infection group (Wilcoxon test p<0.001). Pre-infection and post-infection groups indicate that Losartan was added one hour before (pre-infection) and after the cells were infected (post-infection) with SARS-CoV-2, respectively.

**Fig. 5.**
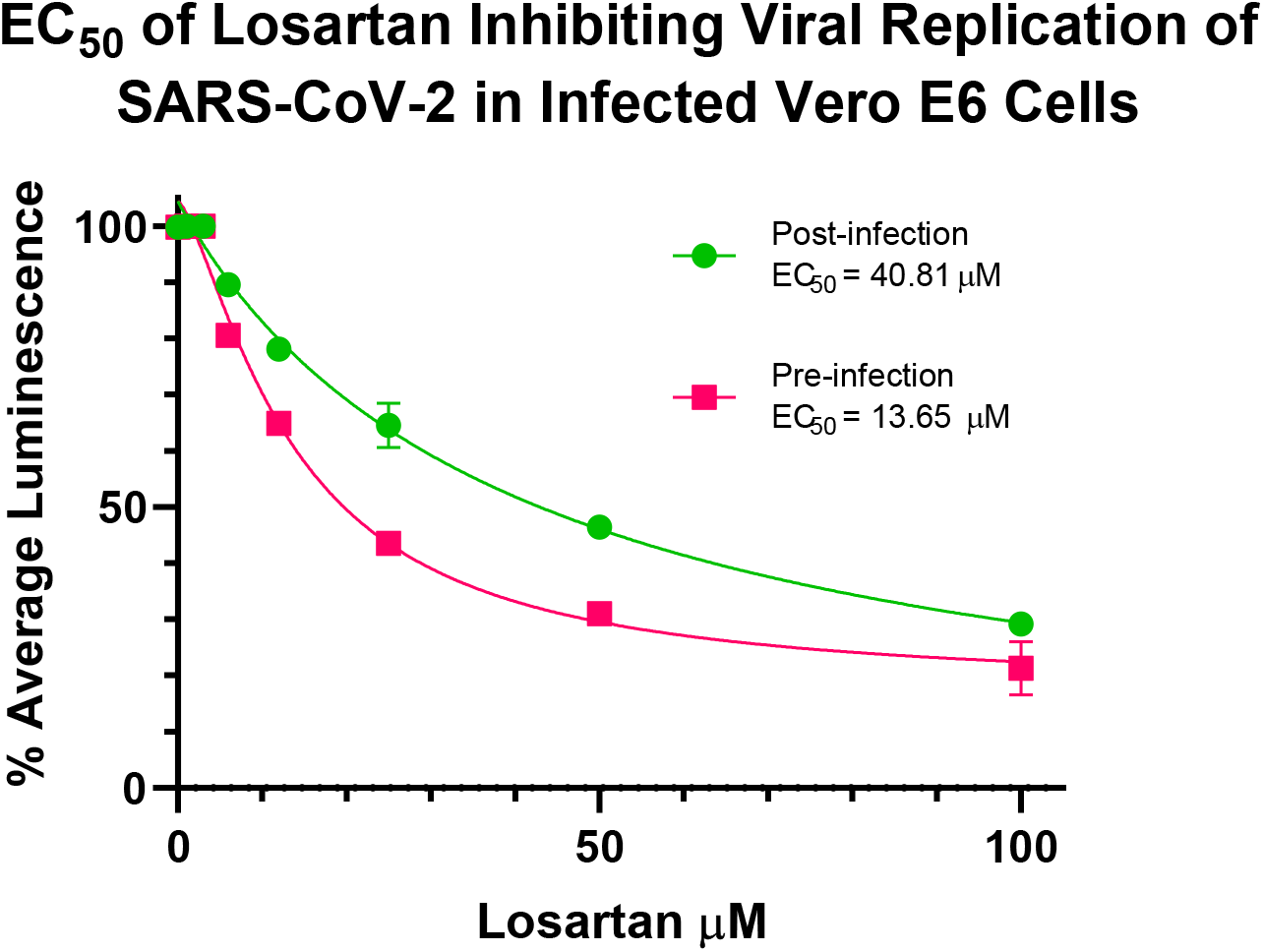
EC_50_ curves in pre- and post-infection groups in 96-well assay. Losartan dose-dependently reduces the production of viral nuclear proteins in Vero E6 cells (near 80%). EC_50_ curves are significantly different in pre- and post-infection groups, significantly (P<0.001).

## Discussion

Replication of SARS-CoV-2 depends on PLpro and Mpro functions. Inhibitors of SARS-CoV PLpro, in several studies, have been shown to be as effective against SARS-CoV-2 PLpro [44, 45]. Although many of the findings were promising, the development in introducing effective drugs is lagging. In 2008, discovery of successful antivirals against SARS-CoV was paused after spread of the virus was contained. Discoveries in disease pathogenesis of SARS-CoV-2 can aid in discovery of novel treatments and drugs. One hypothesis proposes that dysregulation in renin-angiotensin system (RAS) components contribute to the cytokine storm in COVID-19. Downregulation of ACE2 during the entry of the virus into the host cells leads to an increase in angiotensin II/angiotensin(1-7) ratio with proinflammatory, pro-apoptotic, destructive and pro-thrombotic effects [27, 46]. In animal studies, it was shown that ACE2 deficiency in mice and infusion of Ang II in swine resulted in corneal inflammatory response and lung pathology, respectively, resembling exactly the immunopathological changes seen in COVID-19 [47, 48]. In an in-silico study, Losartan was shown to change the tertiary structure of ACE2 in binding with receptor binding domain (RBD) of S protein of the virus as well as inducing modification in conformational shape of SARS-CoV-2 papain-like protease [27]. Accordingly, Losartan may attenuate the inflammatory response and improve organ dysfunctions associated with COVID-19 by rebalancing RAS components and possibly reducing viral affinity for ACE2. This ligand may inhibit the PLpro function as it poses on the same position as the PLpro inhibitor, GRL-0617, does (Fig. 6) [27].

**Fig. 6.**
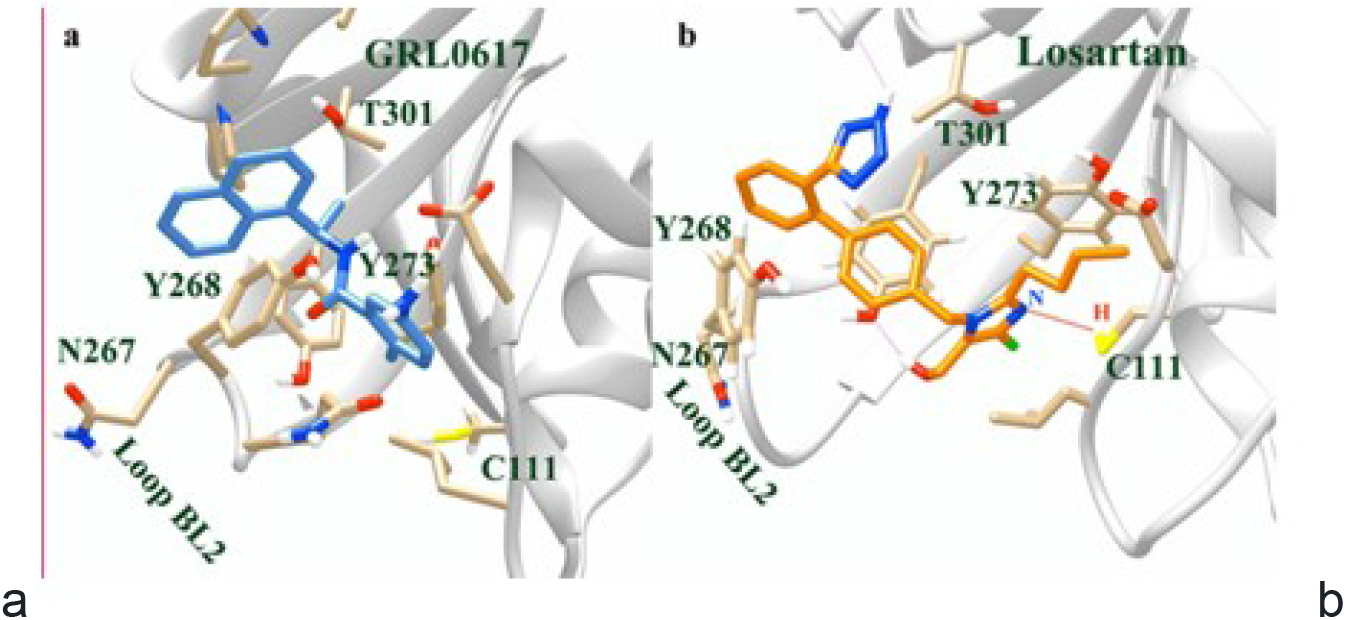
Positioning of GRL-0617 (PLpro inhibitor) and Losartan in the active site of PLpro. (a) X-ray crystallography structure of PLpro and GRL-0617 (PDB ID: 7cmd). (b) Losartan positioning was achieved after 100ns MD simulation. Losartan poses in the same position of the inhibitor (GRL-0617) according to PLpro amino acids in the circumferential area of these two ligands. Hydrogen bonding between nitrogen (N) in Losartan imidazole ring and sulfhydryl group of cysteine (C111) may inhibit initial phase of attacking of cysteine (C111) to the cabonyl group of the substrate in the general cleavage chemical reaction (Fig. 9). Note: X-ray crystallography in contrast to NMR analysis and theoretical modeling cannot resolve hydrogen atoms in most protein crystals found in PDB files [49].

Our in vitro study revealed that Losartan was a weak inhibitor of PLpro in cleavage of peptide-AMC (IC50=1200±61μM). It did not substantially suppress deISGylating ability of PLpro (Fig. 2). It is possible that Losartan interacts with the P3/P4 pocket of PLpro adjacent to its active site and is displaced in the presence of ISG15. This pocket could facilitate cleavage of Ubl substrates by binding the C-terminal of leucine and arginine, of the RLRGG motif, through a protonating reaction in a cleft in the PLpro containing a conserved triad of Histidine(H272)-Cystein(C111)-Aspartic Acid(D286) in papain-like proteases [22, 24, 40]. This is in agreement with the previous bioinformatic findings showing a decrease in the affinity of Losartan for PLpro in the presence of UBl1 (Fig. 7, Fig. 8) [27].

**Fig. 7.**
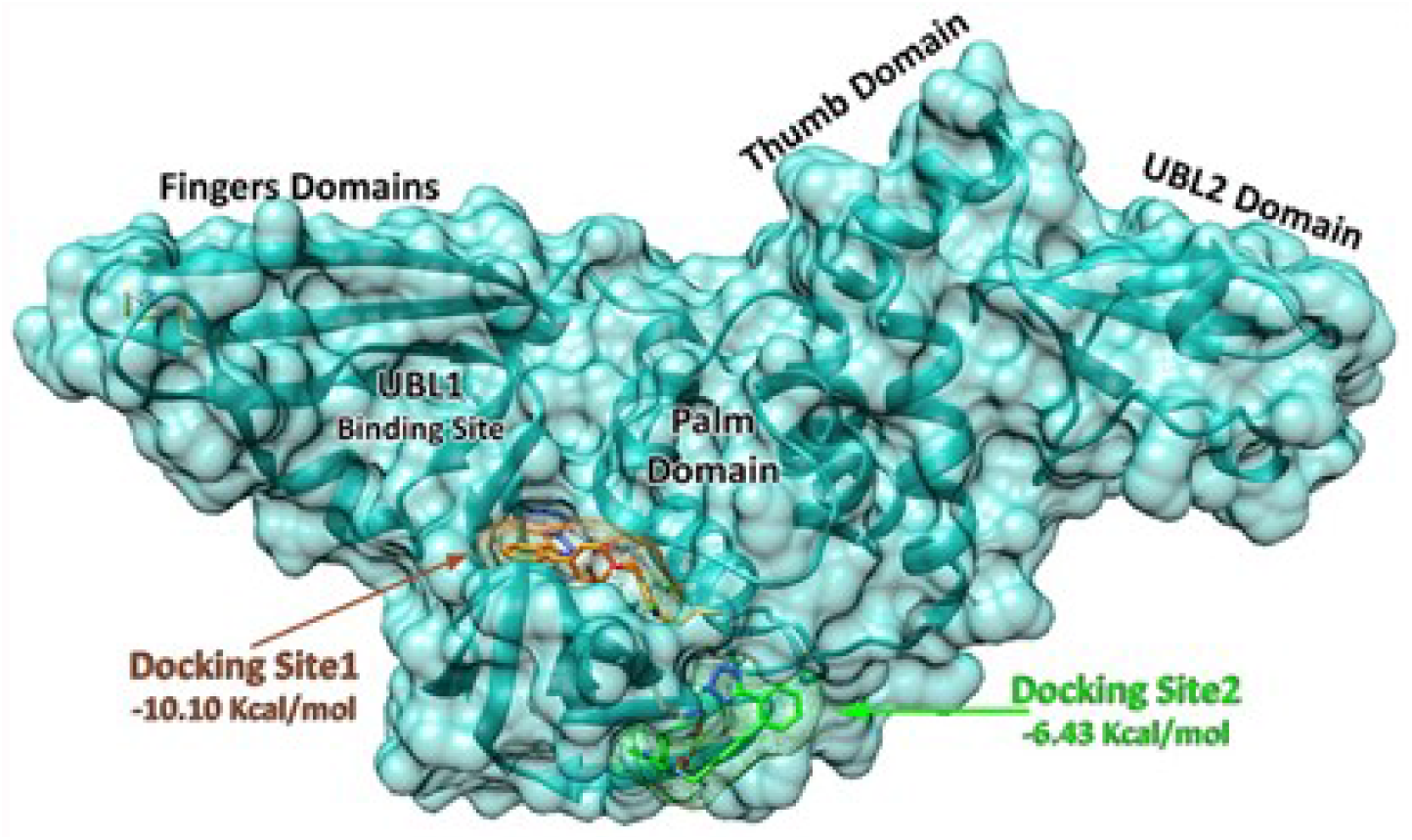
PLpro thumb-palm-finger structure and two positions of Losartan in docking with PLpro. Docking site 1 with - 10.10 Kcal/mol energy is the preferred pose for Losartan in the absence of UBl1. Docking site 2 with −6.43 Kcal/mol energy indicates the preferred pose for Losartan in the presence of UBl1. This demonstrates the decrease in affinity of Losartan in docking with PLpro in the presence of UBl1. This is in concordance with the finding that UBl1 and ISG15 might dislocate Losartan as a competitive inhibitor.

**Fig. 8.**
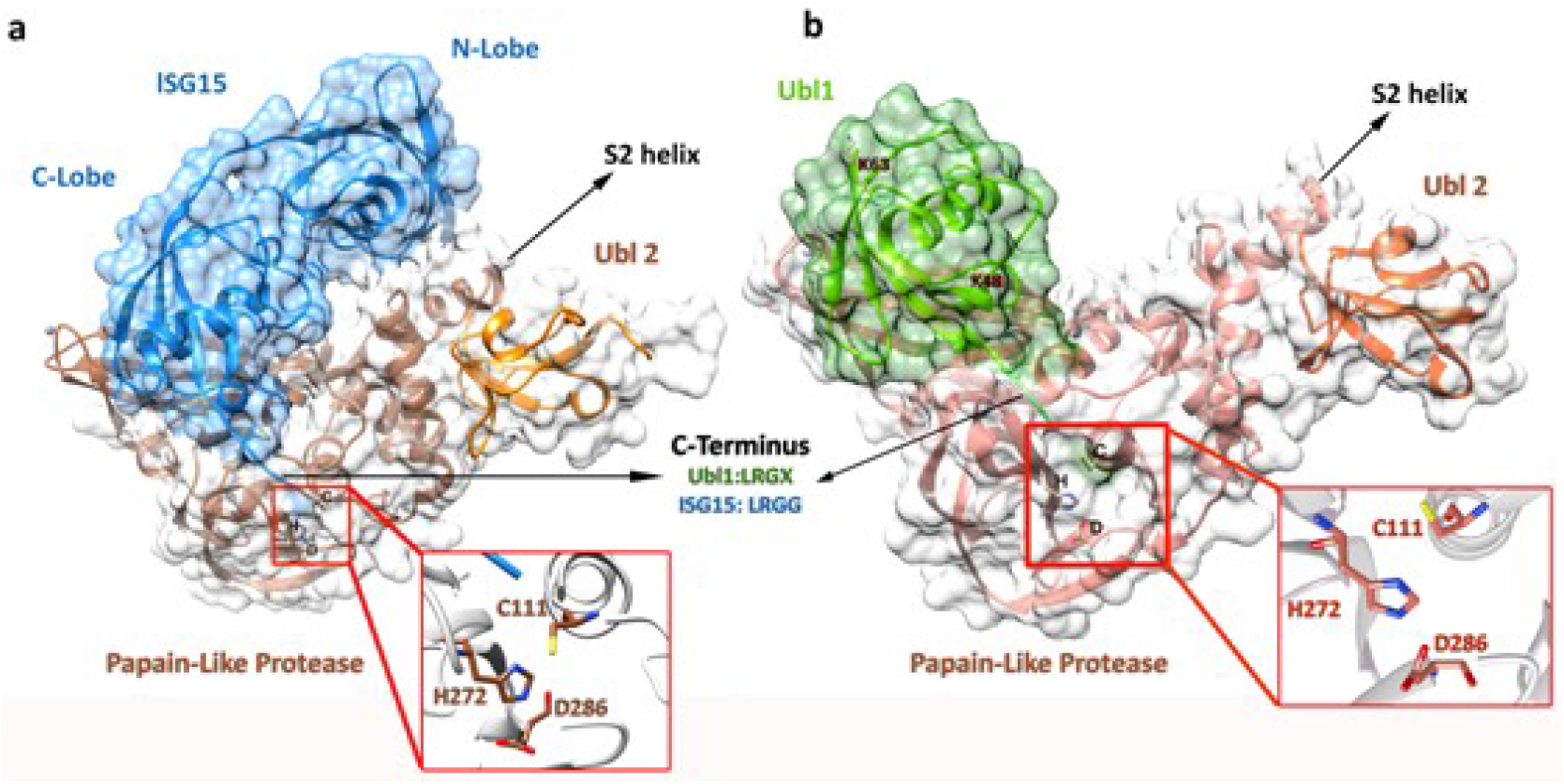
The interaction of SARS-CoV-2 papain-like protease with ISG15 and UBl1 after 100ns MD simulation. The cleavage site of PLpro comprises of a triad of Histidine(H272), Cysteine(C111) and Aspartic acid(D286). C-terminal LXGG of UBl1 and ISG15 are seen in the vicinity of the active site. PLpro crystal structure in interaction with ISG15 (a) was retrieved from protein data bank (PDB ID: 6yva) in which the active site contained Serine (S111) in place of Cysteine (C111) as a point mutation. To emulate the wild structure of PLpro, Serine was replaced by Cysteine and a run of 100ns MD simulation was performed with the new triad.

It has been reported that SARS-CoV-2 PLpro prefers deISGylation 10 times more than deubiquitination [14]. However, our study demonstrated that Losartan in 2.25mM concentration could completely suppress cleavage of monoubiquitin molecules from polyubiquitin chains by PLpro (Fig.1, Fig. 2).

Active (cleavage) site of PLpro with its canonical thumb-palm-finger like structure (Fig. 7), similar to other cysteine proteases in Coronaviruses, comprises two amino acids, histidine and cysteine, to which aspartic acid is added in the case of SARS-CoV and SARS-CoV-2 [22]. The maximum rate of papain-catalyzing activity occurs in pH of 6.2 where both imidazolium ion (ImH^+^) (protonated form of imidazole ring) of histidine and thiolate (deprotonated) form of sulfhydryl group of cysteine are available [50]. Assuming that intracellular pH is near 7 and drops in the context of intracellular ROS overproduction in inflammations like what happens in COVID-19 [27, 51], active site of PLpro finds a favorable milieu to cleave nsps efficiently. As a general catalytic reaction, papain-like protease thiolate ion of cysteine produces a tetrahedral intermediate by attacking the carbonyl group of the substrate which is then released through protonation by ImH^+^ (Fig. 9) [50].

**Fig. 9.**
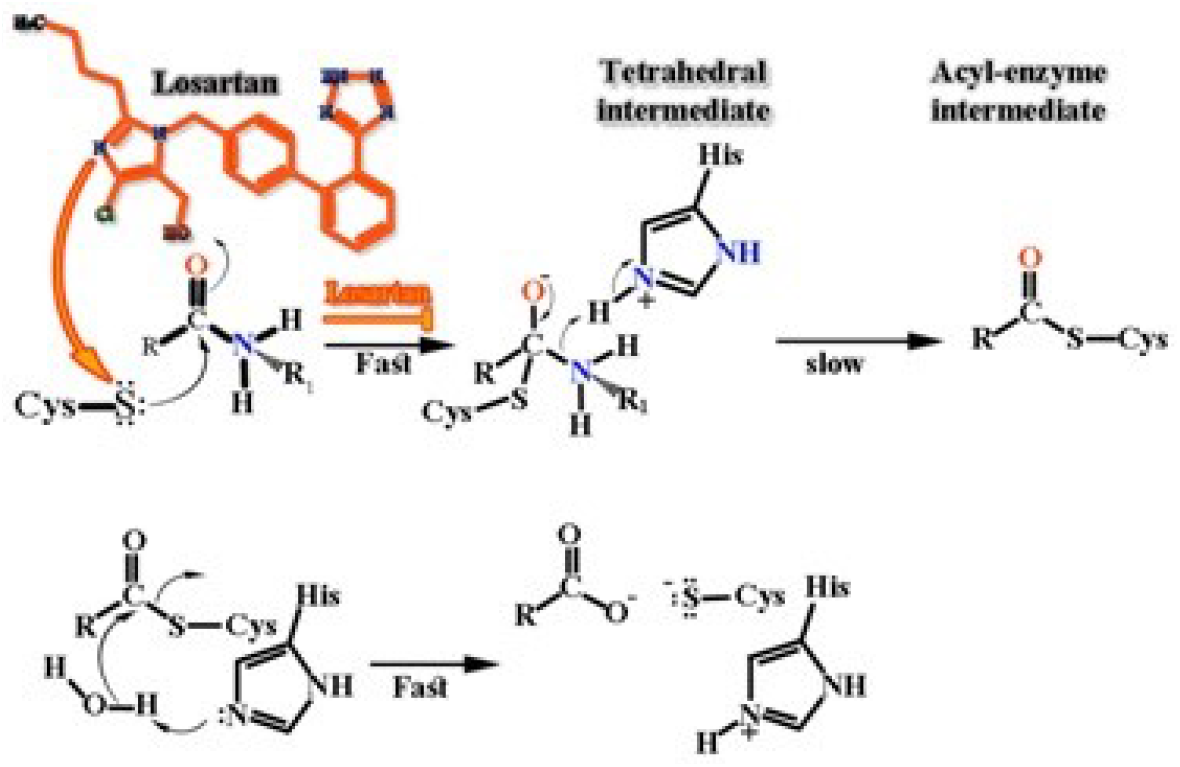
General chemical reaction in cleavage of the carboxy terminal of substrates of PLpro. While cysteine attacks the carbonyl group of a substrate polypeptide and produces an acyl-enzyme intermediate, relaying of proton through imidazole ring of histidine with supporting protonating activity of aspartic acid (D286) in case of PLpro and with the aid of a water molecule in case of Mpro, releases the substrate from cysteine and the triad once again resumes its active configuration. According to our 100ns bioinformatic analysis, nitrogen in imidazole ring of Losartan through protonation of sulfhydryl group of cysteine likely impedes production of tetrahedral intermediate.

A MD analysis [52] revealed that tertiary structure and folding of PLpro in its cleavage cleft changes cysteine (C111) and histidine (H272) to negatively charged deprotonated cysteine and positively charged doubly protonated histidine, respectively. In this context, negatively charged aspartic acid (D286) aids in stabilization of positively charged histidine (H272) [52]. In fact, imidazole ring is a unique structure in biologic reactions, which by possessing two identical nitrogen atoms, it is responsible for relaying protons from one site to another. If two imidazole rings present closely in the path of proton transfer (ImH^+^…Im) an energy barrier may impede the journey of the protons [53]. Accordingly, changes in pH and electrical charge in the catalytic triad of PLpro or presence of any impedance against protonation in the cleavage reaction may distort hydrogen bonds and reduce catalytic efficiency of PLpro in cleavage of nsp1/nsp2, nsp2/nsp3 and nsp3/nsp4 which occurs in c-terminal end of nsp1, nsp2 and nsp3.

Losartan approaches the triad with its imidazole ring which may compete and impede proton transfer in making hydrogen bond with the imidazole ring of histidine (Fig. 10). It is notable that the active site of Mpro of SARS-CoV-2 is composed of a dyad of histidine (H41) and cysteine (C145) to which a water molecule instead of aspartic acid aids in protonation of histidine (H41) [54]. This dyad contributes to cleavage of nsps through the same chemical reaction seen in Figure 9. Also, imidazole ring of Losartan may disrupt the interaction of histidine and cysteine in this protease.

**Fig. 10.**
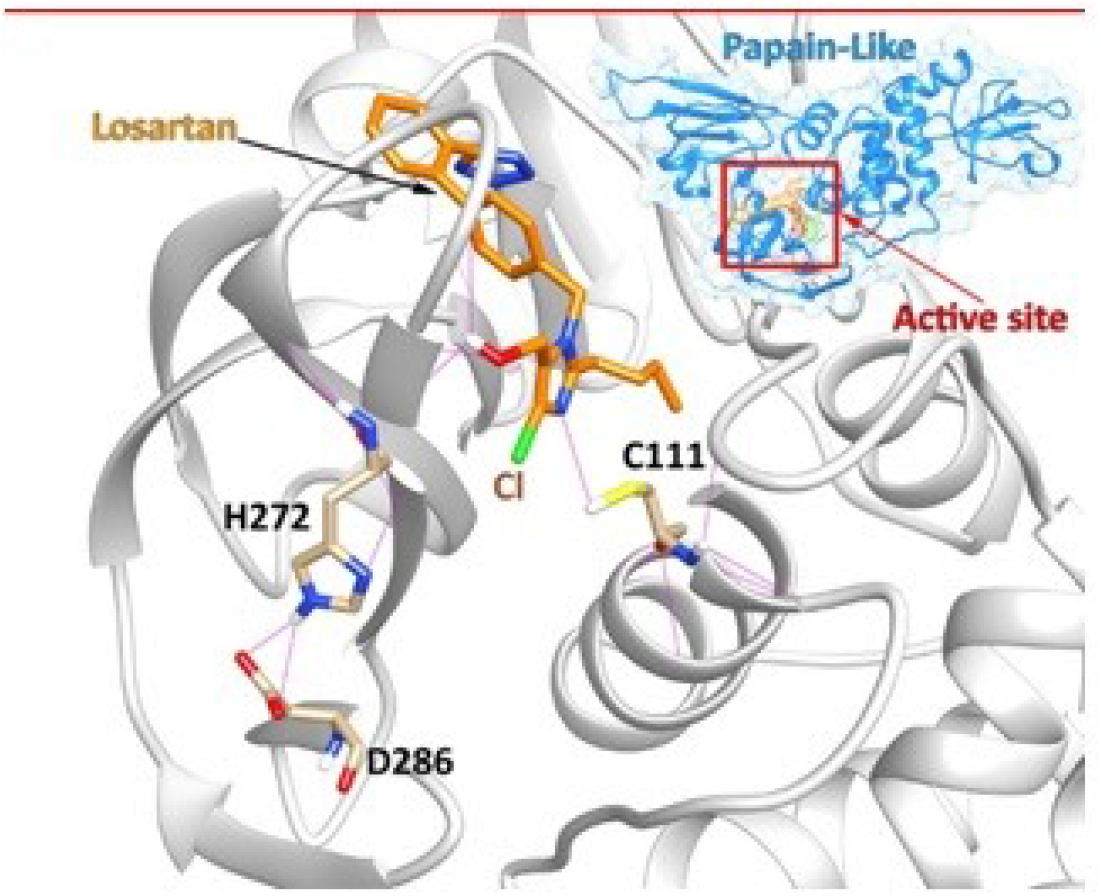
Losartan positioning in the active site of PLpro after 100ns MD simulation. Imidazole ring of Losartan competes with imidazole ring of histidine (H272) for hydrogen bonding with cysteine (C111) in the cleavage triad. Gromacs 2018 package was used for optimization of PLpro (PDB ID: 6xaa) and running 100ns MD simulation through forcefield gromos.

A prior MD analysis [27] demonstrated that Losartan changes the conformational shape of SARS-CoV-2 PLpro. In this study, RMSD graph of 100ns MD simulation of PLpro with and without Losartan shows the changes in the shape of PLpro after 80 ns (Fig. 11) [27].

**Fig. 11.**
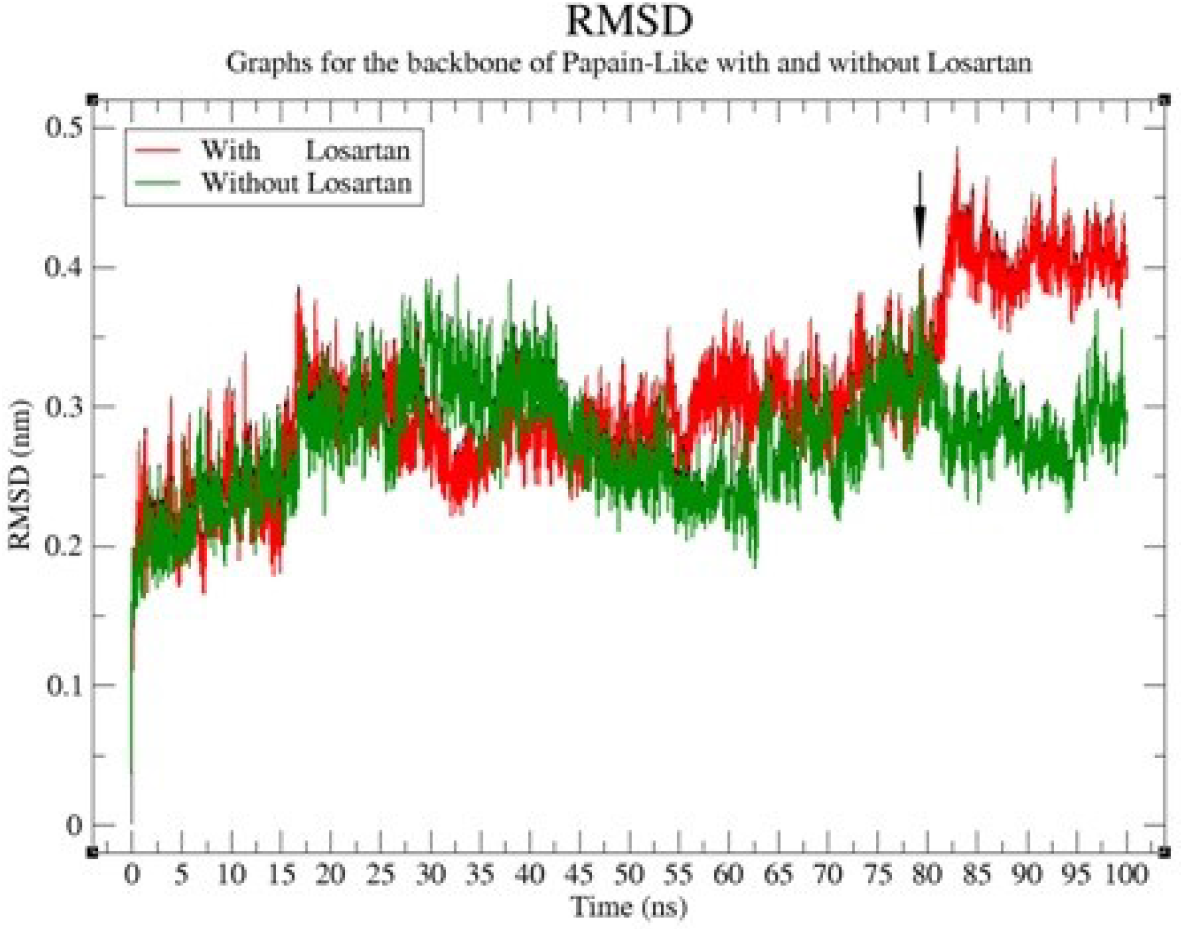
Superimposition of RMSD graphs of 100ns MD simulation of PLpro with (red) and without (green) Losartan. Conformational shape of PLpro changes reasonably after 80ns (black arrow).

Despite the above-mentioned findings in MD simulation, Losartan, in our in vitro study, showed only weak inhibitory effect on PLpro cleavage of monoubiquitin from polyubiquitin chain and only minor inhibition of deISGylation. As to the fact that ubiquitin and ISG15 do not show any homology with at least nsp2 amino acid sequence, it might be likely that cleavage of nsps by PLpro through the general chemical reaction in the active site (Fig. 9) might be associated with different positioning of nsps in PLpro in comparison with ubiquitin and ISG15. Thus, Losartan by penetrating the active site and altering hydrogen bonds in the three effector residues may interfere with cleavage of nsps. Though, this assumption needs further investigation, as an evidence for this concept, our in vitro study showed that 200uM concentration of Losartan could inhibit the cleavage of peptide-AMC containing RLRGG C-terminal sequence up to 35.1% ± 2.7 while at the same concentration, this compound could only inhibit cleavage of ISG15 and UBl1 at 6.9% ± 3.4 and 2.3% ± 1.4, respectively (table 1).

Considering the putative effects of Losartan on Ang II/angiotensin(1-7) ratio, and on ACE2 in interaction with SARS-CoV-2 RBD, PLpro and Mpro, we tested the effect of Losartan on cell destruction and replication by SARS-CoV2 in Vero E6 cell culture.

Vero E6 cells derived from the kidney of African green monkeys, strongly express ACE2 on their apical membranes as observed in certain cells of the respiratory tract [55–57]. Additionally, these cells do not produce type I interferons (IFN type I) allowing SARS-CoV and SARS-CoV-2 replication without being recognized by cell innate immunity [58–63]. According to the results of our Vero E6 cell culture study, which was performed in two-series of triplicated tests, Losartan showed cell protection in both pre- and post-infection groups in a dose dependent manner (p<0.001). Results of clinical trials are expected to be better than *in vitro* results as Vero E6 cells lack interferon producing ability.

Cell destruction in 6-well (number of plaques) and percent of infected cells in 96-well (cells containing nuclear protein) were both reduced significantly (p<0.001) when Losartan was added in increasing concentrations from 0 μM to100 μM by 80% in pre-infection and 70% in post-infection groups (Figures 3 and 4). Based on EC_50_, Losartan when used prior to infecting the cells (pre-infection group) with SARS-CoV-2, reduced NP production by over 50% more than when it is added to the cell culture after infecting the cells with the virus. Therefore, prophylactic use of Losartan may have a significant (p<0.001) clinical benefit if validated by future relevant clinical trials. Additionally, EC_50_ for GRL-0617, a noncovalent inhibitor of PLpro, is reported to be about 27.6 μM [14]. Losartan with EC50 of 7.29 μM in plaque reduction experiment may be more efficient when compared to GRL-0617 in prevention of SARS-CoV2 pathogenesis.

The prominent effect of Losartan in pre-infection group is in agreement with a previously published MD analysis revealing a lower affinity of the virus for ACE2 in the presence of Losartan [27]. Furthermore, the cell surviving potential of Losartan might likely be an evidence supporting the previous bioinformatic finding that this drug could change the conformational shape of PLpro or interfere with the interaction of functional residues (histidine, cysteine) in PLpro or Mpro of the virus [27].

In previous studies, Losartan was described to have anti-inflammatory, anti-apoptotic and anti-thrombotic effects through downregulating Ang II mediated AT1R stimulation [27, 64–71]. Entry of SARS-CoV-2 is associated with downregulation of ACE2 and a subsequent acute rise in Ang II/angiotensin (1-7) ratio with pro-inflammatory, pro-apoptotic, and prothrombotic outcomes [72–75]. In order to control SARS-CoV-2-induced inflammatory responses, many experts have been trying to extend the half-life of ACE2, given the protective role of ACE2 in COVID-19 [76]. As an alternate to restoring ACE2, Losartan, which rebalances the dysregulated RAS in COVID-19, may attenuate ARDS in patients with COVID-19 [27]. Series of unpublished clinical observations revealed the ability of Losartan in suppressing lung inflammation in this disease. Clinical trials are underway to assess efficacy of Losartan in treatment of COVID-19. Data from clinical trial combined with the data from this study may improve the management of this deadly disease.

## Conclusion

COVID-19 has resulted in widespread social, economic, and healthcare crisis. Addition of effective antiviral successful vaccination and antiviral treatment strategy against SARS-CoV-2 would greatly aid in the control of this pandemic. Losartan in this study showed weak inhibitory effect on viral PLpro deubiquitinase and deISGylase properties. This study also showed that Losartan added to the cell culture of Vero E6 cells prior to and after infection of the cells with SARS-CoV2 can inhibit SARS-CoV-2 cell destruction and replication. This, if validated by ongoing clinical trials in different continents, may mean that Losartan has both prophylactic and therapeutic effects in the treatment of COVID-19. Additional experiments are needed to better assess the structural changes in the viral proteins and their biological products when exposed to Losartan. Outcome of these experiments may provide a viable pathway for antiviral design and development. If validated in clinical trials, Losartan with its inhibitory effect on viral replication and its significant benefits in cell protection without any cell toxicity may play an important role in slowing the spread of the disease, medical management of individuals unresponsive to vaccination or new mutations, easing healthcare cost and burden in communities globally.

## Methods

### Chemicals and Reagents

Z-RLRGG-7-amino-4-methyl-courmarin (peptide-AMC) was purchased from Bachem. Ubiquitin−7-amino-4-methylcourmarin (Ub-AMC) was purchased from Boston Biochem; human ISG15−7-amino-4-methylcourmarin (ISG15-AMC) was purchased from Boston Biochem. Lys48 linked tetra-Ub were obtained from Boston Biochem; DL-dithiothroitol (DTT) was purchased from GoldBio, and isopropyl-β-D-thiogalactopyranoside (IPTG) was purchased from GoldBio. 4-(2-Hydroxyethyl)-1-piperazineethanesulfonic acid (HEPES) was purchased from Fisher BioReagents. Imidazole was purchased from Acros Organics; tris(hydroxymethyl)aminomethane (Tris) was purchased from Fisher Scientific. Losartan and Sodium chloride (NaCl) were purchased from Fisher Chemical, and bovine serum albumin (BSA) was purchased from Sigma Life Science.

### Determination of IC_50_ Values

IC_50_ assays were performed using methods previously described for peptide-AMC cleavage experiments [45]. All assays were run using Corning Costar half-volume 96-well plates containing AMC buffer (100mM NaCl, 50mM HEPES [pH = 7.5], 0.01mg/mL bovine serum albumin (BSA), and 5mM DTT) to a final volume of 50μL and performed in triplicate. The CLAIROstar plate reader (BMG Lab Tech, Inc.) was used to measure the fluorescence of the AMC cleavage, and the data was analyzed using MARS (BMG Lab Tech, Inc.). Assays using peptide-AMC substrate contained 1μM SARS-CoV-2 PLpro and 50μM peptide-AMC in 98% AMC buffer/2% DMSO. Reactions were performed in triplicate with inhibitor concentrations ranging from 390nM to 2.25mM. Further assays utilizing ISG15-AMC and Ub-AMC contained 1nM PLpro with 1μM substrate or 25nM PLpro with 2μM substrate respectively. IC_50_ calculations were performed using Prism8 from GraphPad.

### Inhibition of Poly-Ub and proISG15 Cleavage

Lys48 linked tetra-Ub obtained from Boston Biochem was incubated at 10.5μM with 23nM SARS-CoV-2 PLpro and 2.25mM Losartan. Reactions were performed in 97.75% AMC buffer/2.25% DMSO at a volume of 80μL and a temperature of 37°C. 10μL samples were taken at the indicated time points and heat-shocked at 98°C for 5 min. SDS-PAGE analysis was performed using Mini-PROTEAN TGX and Coomassie blue. Utilizing similar parameters 10μM human proISG15 was incubated with 20nM PLpro and 2mM Losartan. Reactions were performed in 98% AMC buffer/2% DMSO at a volume of 90μL. 10μL samples were taken at the indicated time points and heat-shocked at 98°C for 5 min. SDS-PAGE analysis was performed using Mini-PROTEAN TGX Stain-Free.

### 6-Well and 96-Well Losartan Antiviral Assays

In the 6-well setting, plaques represent the destructed cells, while in the 96-well test the percentage of Relative Light Unit (RLU) represents detection of viral nuclear protein (NP) in the infected cells.

### SARS-CoV-2 Post-infection Cells and Losartan Antiviral Assays

SARS-CoV-2 (2019-nCoV/USA-WA1/2020; MN985325.1) was received from BEI resources and propagated in Vero E6. Infections were done at a multiplicity of infection (MOI) of 0.1 in serum-free in Dulbecco’s minimal essential medium (DMEM) for 1h after which the virus-containing media was decanted and replaced with DMEM supplemented with 1% heat-inactivated fetal bovine serum [77]. The virus was propagated for 72 hours before it was harvested and the titer determined by plaque assay on Vero E6 cells [78]. The viral plaques were counted and the titer was determined as PFU/ml. The Vero cells were plated at 5 × 10^5^ cells/well in a 6-well plate and incubated overnight at 37°C. The following day the Losartan was prepared into the following concentrations/well in a separate plate; 100 μM, 50 μM, 25 μM, 12 μM, 6 μM, 3 μM and 1 μM. The cells were washed once with PBS once and then infected at a multiplicity of infection (MOI) of 0.01 for 1h after which the virus containing media was removed and the Losartan concentrations were added to the cells with 3 mL overlay and incubated for 96 hours at 37°C at 5% CO_2_. The cells were then fixed and stained with crystal violet to determine plaque numbers. These were all done in triplicate and the calculations were performed using Prism8 from GraphPad. In the 96-well assay, 200 μL of the media containing the Losartan concentrates and the virus was incubated for 96 hours, in triplicate. This mixture was fixed, stained for Viral nuclear protein (NP), and assay on Cellomics arrays was performed. Relative Light Unit (RLU) detecting viral NP correlates with replication the infected cells [79]. Average percentage responder intensity of the fluorescent channel from NPs of the virus was measured and the calculations were performed using Prism8 from GraphPad.

### SARS-CoV-2 Pre-infection Cells and Losartan Antiviral Assays

SARS-CoV-2 (2019-nCoV/USA-WA1/2020; MN985325.1) was received from BEI resources and propagated in Vero E6. Infections were done at a multiplicity of infection (MOI) of 0.1 in serum-free in Dulbecco’s minimal essential medium (DMEM) for 1h after which the virus-containing media was decanted and replaced with DMEM supplemented with 1% heat-inactivated fetal bovine serum [77]. The virus was propagated for 72 hours before it was harvested and the titer determined by plaque assay on Vero E6 cells [78]. The viral plaques were counted and the titer was determined as PFU/ml. The Vero cells were plated at 5 × 10^5^ cells/well in a 6-well plate and incubated overnight at 37°C. The cells were washed once with PBS once. Losartan was prepared into the following concentrations/well in a separate plate; 100 μM, 50 μM, 25 μM, 12 μM, 6 μM, 3 μM and 1 μM. For a total of 3 mL final volume per well, Losartan concentrate were added to the Vero E6 wells and incubated for one hour. Viral dilution was then made at a multiplicity of infection (MOI) of 0.01 after which the virus containing media was removed and the Losartan concentrations were added to the cells with 3 mL overlay and incubated for 96 hours at 37°C at 5% CO_2_. The cells were then fixed and stained with crystal violet to determine plaque numbers. These were all done in triplicate and the calculations were performed using Prism8 from GraphPad. In the 96-well assay, 200 μL of the media containing the Losartan concentrates and the virus was incubated, in triplicate. This mixture was fixed, stained for Viral NP, and assay on Cellomics arrays was performed. Relative Light Unit (RLU) detecting viral nuclear protein (NP) correlates with viral replication in the infected cells [79]. Average percentage responder intensity of the fluorescent channel from NPs of the virus was measured and the calculations were performed using Prism8 from GraphPad.

## Author Contributions

The manuscript was written through the contributions of all authors. All authors have approved the final version of the manuscript.

## Acknowledgments

We thank our families and friends for their support which provided us with the time to pursue this endeavor while they endured the challenges that all families have encountered in this pandemic.

The humanitarian research in this article was made possible by the University of Georgia College of Pharmacy, Office of Research, College of Veterinary Medicine, and the University of Georgia’s Center for Drug Discovery (S.D.P).

We thank Derren Barken, PhD for helping with verification of our statistical analysis of the data presented in this research.

We thank California Medical Innovations Institute for their reception and encouragement.

## Abbreviations

ACE: Angiotensin-Converting Enzyme
ACE2: Angiotensin-Converting Enzyme 2
AMC: C-terminal 7-amido-4-methylcoumarin
Ang I: Angiotensin I
Ang II: Angiotensin II
Ang (1-7): Angiotensin 1-7
Ang (1-9): Angiotensin 1-9
ARB: angiotensin receptor blockers
ARDS: acute respiratory distress syndrome
AT1R: Angiotensin II type 1 receptor
AT2R: Angiotensin II type 2 receptor
BSA: bovine serum albumin
CoV: coronavirus
COVID-19: coronavirus disease 2019;
DMSO: dimethyl sulfoxide
DUB: deubiquitinase
GRL-0617: 5-amino-2-methyl-Nbenzamide
IL-6: interleukin-6
ISG15: interferon-stimulated gene product 15
MD: molecular dynamic
MERS-CoV: Middle East Respiratory Syndrome
NP: nuclear protein
NSP: nonstructural proteins
ORF1a/ORF1b: open reading frames
PLP: papain-like protease
PLpro: papain-like protease
proISG15: precursor ISG15
PRR: Pattern-Recognition Receptor
RAS: renin-angiotensin system
RBD: receptor binding domain
SARS-CoV-2: severe acute respiratory syndrome coronavirus 2
TLR: Toll-like receptor
Tris: tris(hydroxymethyl)-aminomethane
Ub: ubiquitin
poly ub: polyubiquitin
UIM: ubiquitin interacting motif

